# A common goodness-of-fit framework for neural population models using marked point process time-rescaling

**DOI:** 10.1101/265850

**Authors:** Long Tao, Karoline E. Weber, Kensuke Arai, Uri T. Eden

## Abstract

A critical component of any statistical modeling procedure is the ability to assess the goodness-of-fit between a model and observed data. For neural spike train models of individual neurons, many goodness-of-fit measures rely on the time-rescaling theorem to assess the statistical properties of rescaled spike times. Recently, there has been increasing interest in statistical models that describe the simultaneous spiking activity of neuron populations, either in a single brain region or across brain regions. Classically, such models have used spike sorted data to describe relationships between the identified neurons, but more recently clusterless modeling methods have been used to describe population activity using a single model. Here we develop a generalization of the time-rescaling theorem that enables comprehensive goodness-of-fit analysis for either of these classes of population models. We use the theory of marked point processes to model population spiking activity, and show that under the correct model, each spike can be rescaled individually to generate a uniformly distributed set of events in time and the space of spike marks. After rescaling, multiple well-established goodness-of-fit procedures and statistical tests are available. We demonstrate the application of these methods both to simulated data and real population spiking in rat hippocampus.

## 1 Introduction

Statistical models have proven to be a powerful approach to capturing the coding properties of neural systems (Brown et al, 2004; Kass et al, 2005; Paninski et al, 2007; Kass et al, 2014). In addition to describing the associations between spiking activity and the biological and behavioral signals being represented, they also provide tools for model assessment and refinement. As electrophysiological experiments have become more sophisticated, incorporating simultaneous spiking data from more neurons across multiple brain areas, the focus of neural data analysis problems has begun to shift from ones that attempt to understand the tuning properties of individual neurons to ones that attempt to capture the combined structure of activity from neural populations (Georgopoulos et al, 1986; Wu et al, 2002; Pillow et al, 2008; Paninski et al, 2009; Shanechi et al, 2012). This shift has generated a need for statistical modeling and goodness-of-fit tools that can address neural coding problems at the population level.

Considering spikes as localized events in time, which are most appropriately described using the theory of point processes, has led to a class of statistical models that has been highly successful at capturing the coding properties and dynamics of individual neurons (Kass and Ventura, 2001; Truccolo et al, 2005; Pillow et al, 2008). The traditional neural point process modeling framework relates the spiking activity of isolated or sorted neurons to their own recent spiking history, that of other neurons in its network, and to the behavioral and biological signals to which the neurons respond (Brown et al, 2002; Smith and Brown, 2003; Truccolo et al, 2005; Deng et al, 2013; Arai and Kass, 2017). Notable examples include modeling of spatial coding and movement trajectories using firing in the CA1 region in the rat hippocampus (Brown et al, 1998; Huang et al, 2009; Eden et al, 2018), as well as the neural decoding of hand velocities and collective dynamics in the primary motor cortex (Georgopoulos et al, 1986; Eden et al, 2004; Brockwell et al, 2004; Srinivasan et al, 2006). Relating population neural activity to behavior may be improved if instead of using spikes sorted according to neural identity, sorting is skipped entirely, and a joint model of behavior and features of unsorted spike waveforms across the neural population is built directly (Kloosterman et al, 2014; Deng et al, 2015; Sodkomkham et al, 2016).

Models of this type can be described using the theory of marked point process models (Daley and Vere-Jones, 2003). In this case, the mark could be the full spike waveform, but is often taken instead to be some feature or low dimensional set of features related to the waveform, such as amplitude or half-width. Marked point process models can also be used to describe spiking activity from populations of sorted spikes, where the mark is often a discrete label indicating into which cluster each spike was sorted. Due to the generality of this class of marked point process models and its ability to model both sorted and unsorted population spiking data, it is of great importance that a corresponding set of tools for model assessment and validity, commonly referred to as goodness-of-fit, be developed to enable accurate model assessment and interpretation of the resulting fits. When properly developed and implemented, these types of model assessment metrics are helpful for determining whether a model accurately reflects the structure of a neural representation and whether the representation remains stable in the face of experimental dynamics. They can also provide a way to further refine a given model and understand the specific ways in which it may be underperforming or lacking fit.

Multiple goodness-of-fit tools have been established for point process models of individual neurons. Notably, many of these methods are based on a fundamental theoretical result known as the time-rescaling theorem (Papangelou, 1972; Brown et al, 2002), which indicates that any point process representing a neural spike train can be rescaled based on its instantaneous spiking intensity so that it becomes a simple Poisson process with a constant spike rate. In terms of model assessment, this means that for any proposed neural spiking model, we can rescale the observed spikes according to that model and assess the goodness-of-fit between the rescaled spiking and the known properties of Poisson processes. Notably, researchers often use Kolmogorov-Smirnov (KS) plots, which compare the empirical distribution of the rescaled interspike intervals to the distribution of interspike intervals expected from a Poisson process. This is one of a range of goodness-of-fit tools made available through the time-rescaling approach.

However, with the expanding development of these new marked point process models for population data, there is a need for a corresponding development of appropriate goodness-of-fit tools that can be applied generally to these models. Gerhard et al (2011) describe an approach based on time-rescaling multiple univariate point processes. Vere-Jones and Schoenberg (2004) prove the general time-rescaling theorem for marked-point processes, but do not develop its use for goodness-of-fit over a fixed observation interval. In this paper, we describe a new methodology that extends these approaches, based on a generalization of the time-rescaling theorem to marked point processes. We provide a heuristic proof of the theorem, and illustrate the method with simulated and real data from population spiking.

The key idea behind this generalization is to consider a marked point process model as providing a description of the spiking intensity about a neighborhood of any mark value, and to rescale each observed spike individually, based on its mark. The marked point process time-rescaling theorem then indicates that the resulting rescaled marked point process has spikes that are uniformly distributed in time and mark space, in a region that is defined by rescaling the observation interval, [0,T] across all marks. Therefore, assessment of the marked point process models can be performed using goodness-of-fit techniques for uniformity of the spikes. Additionally, by taking the superposition of the rescaled spikes over all marks we obtain a univariate point process in time and a rescaled intensity. If the original marked point process model is correct, the resulting process will be an inhomogeneous Poisson process with the given intensity, allowing for the use of standard point process goodness-of-fit tools such as KS plots. In fact, this procedure allows for an extensive array of goodness-of-fit techniques that aid in model assessment and refinement for the inhomogeneous Poisson process.

Section 2 will provide a brief summary of point process modeling methods and the time-rescaling theorem in a single dimension for general univariate point processes, followed by a description of the approach for modeling neural populations as marked point processes. We will then describe a generalization of the time-rescaling theorem for these models, and provide a heuristic proof of the theorem. In sections 3 and 4, we illustrate our model assessment method by simulation as well as a real-data application, respectively.

## 2 Methods for goodness-of-fit based on the time-rescaling theorem

### 2.1 The conditional intensity function and the time-rescaling theorem for univariate point processes

Define an observation interval [0, *T*] and let 0 ≤ *s*_1_ < *s*_2_ <,…,< *s_n−1_* < *s_n_* ≤ *T* be a set of event (spike) times. Let *N*(*t*) be the number of spikes up to time *t*, which will increase by 1 at times when a spike occurs and will remain constant otherwise. Any point process *N*(*t*) describing neural spiking can be fully characterized by its conditional intensity function (Daley and Vere-Jones, 2003)

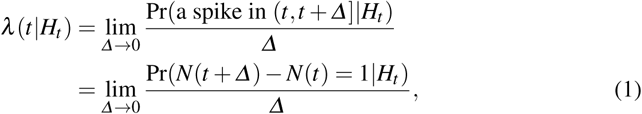

where *H_t_* = {0 ≤ *s*_1_ < *s*_2_ <,…,< *s*_*N*(*t*)_ ≤ *t*} is the history of spiking activity up to time *t*. The conditional intensity function expresses the instantaneous likelihood of observing a spike at time *t*, and implicitly defines a complete probability model for the point process. It therefore serves as the fundamental building block for constructing the likelihoods and probability distributions needed for the point process data analysis.

The basic idea of the time-rescaling theorem is to transform a general temporal point process to a constant-intensity Poisson process by rescaling the spike times.

**Theorem 1 (time-rescaling theorem)**

For a given point process *N*(*t*) with conditional intensity function *λ*(*t*|*H_t_*) with event (spike) times 0 ≤ *s*_1_ < *s*_2_ <,…,< *s*_*N*(*T*)_ ≤ *T* in an observation interval [0, *T*], let

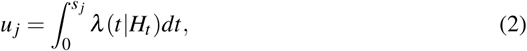

for *j* = 1,…,*N*(*T*). Then *u_j_* are the spike times of a homogeneous Poisson process with unit intensity rate, called the rescaled spike times.

Note that *u_j_*, *j* = 1,…,*N*(*T*), will be independent, identically uniformly distributed on the observation interval 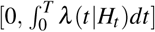 (Ross, 1996). Once a point process model is fitted, we can integrate the estimated conditional intensity between the observed spike times *s_j_*−1 to *s_j_* to get a set of rescaled interspike intervals, 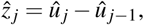 which should be independent, and follow an exponential distribution with rate equal to 1 if the fitted model is correct. This allows for the application of well studied goodness-of-fit methods for assessing models of independent, identically distributed data. For example, the Kolmogorov-Smirnov (KS) plot, which plots an empirical distribution from data against a model distribution, can be used to compare the rescaled interspike intervals to the exponential. Similarly an autocorrelation analysis of the rescaled interspike intervals should show no significant structure at any lag if the estimated conditional intensity from the fitted model accurately describes the spiking observations (Brown et al, 2002; Truccolo et al, 2005).

### 2.2 The joint mark intensity function and the general time-rescaling theorem for marked point processes

We describe spike data from a neural population using a combination of the spike time, and another variable, **m**, called the mark, which can provide information about the spike waveform or the identity of the neuron to which that spike is associated (Kloosterman et al, 2014; Deng et al, 2015). This mark may be discrete (e.g. Neuron 1 vs Neuron 2) or continuous (e.g. spike amplitude); it may be univariate, a vector (e.g. spike amplitude from each channel in a tetrode), or even a function (e.g. a continuous waveform function). The population spiking activity is then given by the set of observations (*s*_1_, **m**_1_), (*s*_2_, **m**_2_),…, (*s_n_*, **m**_*n*_).

A marked point process is completely defined by its joint mark intensity function such that:

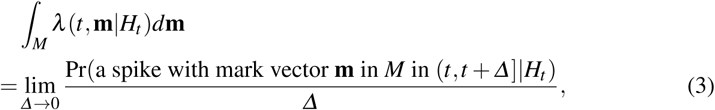

where *M* is a subset (Borel set) of the mark space 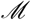 and *H_t_* is the history of spiking activity, including all the marks, up to time *t*. Here *λ*(*t*, **m**|*H_t_*) characterizes the instantaneous likelihood of observing a spike with mark **m** at time *t*. For fixed value m and *t*, *λ*(*t*, **m**|*H_t_*) may depend on the past history of spikes with similar marks (corresponding to the intrinsic history dependence of each neuron), on the history of spikes with dissimilar marks (corresponding to functional connectivity between neurons), and on the extrinsic covariates that the neural population is encoding (for example, place and movement coding in rat hippocampus).

Taking an integral of *λ*(*t*, **m**|*H_t_*) over the entire mark space 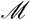,

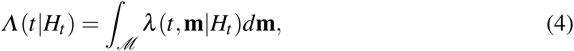

gives the conditional intensity of observing a spike at time *t* regardless of the mark value. *Λ*(*t*|*H_t_*) is often called the ground intensity of the marked point process (Daley and Vere-Jones, 2003).

Marked point process modeling has been successfully applied to multi-unit spiking data (Ba et al, 2014; Kloosterman et al, 2014; Deng et al, 2015). While some theoretical results related to time-rescaling of the marked point processes have been developed (Vere-Jones and Schoenberg, 2004), a complete goodness-of-fit paradigm for population spiking models over fixed observation intervals has yet to be established. Here, we present a general time-rescaling theorem for marked point processes observed on a finite observation interval [0, *T*], with marks that could be either continuous or discrete.

**Theorem 2 (General time-rescaling theorem)**

For a marked point process with observed marks 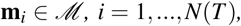 associated with the spike times 0 ≤ *s*_1_ <,…,< *s*_*N*(*T*)_ ≤ *T* and with joint mark intensity function *λ*(*t*, **m**|*H_t_*). Let

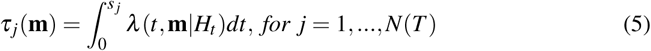

be a set of rescaled spike times, let

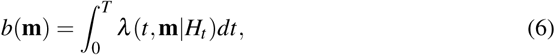

be a mark dependent boundary based on the rescaled value of *T* for each mark, and let *R* = {(*τ*, **m**): 0 ≤ *τ* ≤ *b*(**m**)} be a stochastic region defined by this boundary. Then the joint distribution of 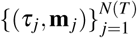 and the number of spikes in region R is equal to that of a homogeneous marked Poisson process with constant mark intensity equal to 1. Therefore, conditional on the boundary *b*(**m**), each (unordered) spike is independently, uniformly distributed in the region *R.*

A heuristic proof of this theorem arises from a simple change of variables. Consider the joint probability distribution of all of the spike times and marks, which is given by the product of the joint mark intensity function, *λ*(*s_j_*, **m**_*j*_|*H_s_j__*), at the spike locations and the exponential of the negative integral of *λ*(*t*, **m**|*H_t_*) over the whole time-mark space:

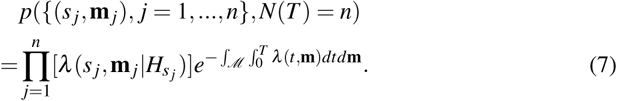

Note that these marked spikes completely specify the joint mark intensity (which is history dependent) everywhere in the observation interval and therefore also specify the extent of the stochastic region *R*. By the multivariate change-of-variables formula (Port, 1994), the joint distribution of the rescaled times and marks is given by the expression:

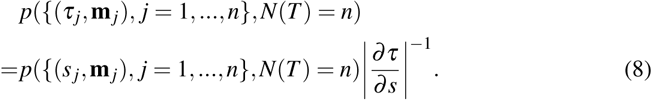

The elements of 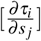 are equal to *λ*(*s_j_*, **m**_*j*_|*H_s_j__*) if *i* = *j*, and are 0 if *i* < *j*. Therefore 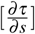 is a lower triangular matrix, and its determinant is given by the product of its diagonal terms, 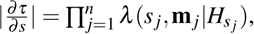 so that

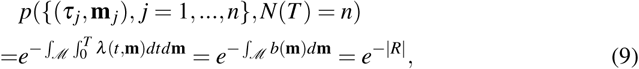

where *|R|* is the volume of region *R*. This is equivalent to the joint distribution of a marked point process with constant unit joint mark intensity over the region *R*.

We can further conclude that the number of spikes in region *R* follows a Poisson distribution with mean equal to *|R|*. Thus the conditional joint distribution of rescaled spike times given that there are *n* spikes in the region *R* is

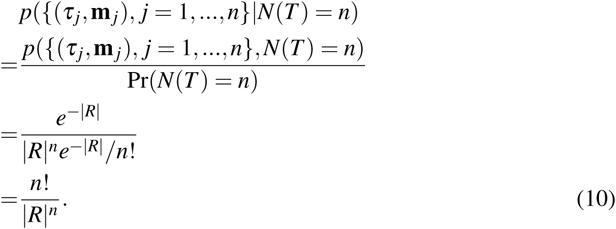

This is exactly the joint density function of a temporally ordered set of independent uniformly distributed events in the rescaled stochastic region *R*.

Here, we presented a heuristic proof of the marked point process time-rescaling theorem based on a change-of-variables argument with the intension of providing intuition about the effect of rescaling. A complete proof requires a few additional details to ensure that the resulting process is well behaved, and more technical proofs are available in the literature (Meyer, 1971; Brown and Nair, 1988; Vere-Jones and Schoenberg, 2004).

Based on the time-rescaling theorem result above, we can also derive the spike rate for the ground process of all the rescaled spikes across all marks.

#### Corollary 1

For a rescaled, marked point process with unit joint intensity function in region *R* as defined above, the (rescaled) spike times will be an inhomogeneous Poisson process with conditional intensity given by

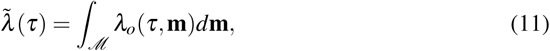

where

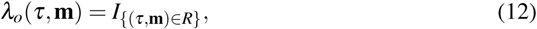

is the indicator function that specifies whether the point (*τ*, **m**) is in the region *R* or not.

### 2.3 Assessing model goodness-of-fit using the general time-rescaling theorem

The marked point process time-rescaling theorem establishes the joint distribution of the rescaled spikes under the assumption that the joint mark intensity model is correct. Therefore, the problem of assessing the goodness-of-fit of any proposed model can be reduced to the simpler problem of determining whether the distribution of the rescaled spike times and marks are consistent with a unit-rate marked Poisson process, or equivalently, whether the spikes occur uniformly over the region *R*.

There are a variety of well studied approaches for assessing goodness-of-fit based on this rescaled process. These multiple methods are complimentary in that one method may detect lack of fit due to particular structure in the data that may not be detected by another method. A number of these are discussed in the discussion section, but here we focus on two relatively simple approaches that are easy to interpret and highlight multiple ways in which the model may fit the data well or poorly.

The first approach is based on Peason’s chi-square statistic. To implement this, we divide the region *R* into *M* smaller subregions, *R_i_*, each with volume |*R_i_*|, and count the number of rescaled spikes, *r_i_*, in each of these subregions. The test statistic is

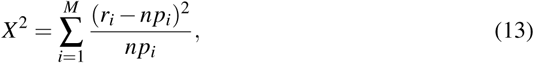

where 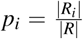 and *n* is the total number of points in *R*. We select the subregions such that *np_i_* is sufficiently large (say, above 5) in each. If our marked point process model is correct and the rescaled spikes are uniform in this region, then *X*^2^ will follow a chi-square distribution with *M* − 1 degrees of freedom. We will reject the null hypothesis that the points are uniformly distributed in region 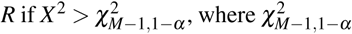 is the critical value of the chi-square distribution with *M* − 1 degrees at a level of significance *α*.

Another approach for assessing the goodness-of-fit for the rescaled process is based on the Kolmogorov-Smirnov (KS) plot. For a univariate (unmarked) point process, if the model is correct, the rescaled process should be a homogeneous Poisson process with interspike intervals that have independent exponential distributions with mean 1. A KS plot then simply plots the empirical cumulative distribution function (CDF) of the rescaled times against the model CDF of an exponential distribution to visualize the deviation from the 45 degree line (Johnson and Kotz, 1970). For a marked point process, the set of rescaled spike times (ignoring the mark values) should be an inhomogeneous Poisson process with rate 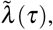 as discussed in the previous section. We can therefore rescale this process one more time, based on the univariate time-rescaling theorem, construct KS plots, and make inferences from them.

We will demonstrate the time-rescaling theorem as well as these two goodness-of-fit approaches to simulated data in section 3, and to real neural population spiking data recorded from a rat performing a memory-guided spatial navigation task in section 4.

## 3 Simulation study

We developed a couple of simple simulation examples to demonstrate the process of using this general time-rescaling approach on spike train data, both for models of sorted spikes and for clusterless models of population spiking.

### 3.1 Simulation study 1

The first simulation scenario comprises two neurons with spiking tuned to a single covariate, *x_t_*, with coordinated, history dependent firing and overlapping mark distributions. We can think of *x_t_* as a one-dimensional position variable, and our neurons as place cells with distinct place fields. Each neuron has a history dependent structure leading to a brief refractory period, and neuron 2 has an excitatory influence on neuron 1 at a lag of 10 time steps.

The position variable, *x_t_*, is modeled as a stationary autoregressive (AR(1)) process. Mathematically, we define the state update equation for *x_t_* as:

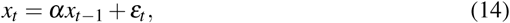

where *α* = 0.98 and *ε_t_* is a zero mean white noise process, with standard deviation 0.3. The top panel of Fig. 1 shows a realization of *x_t_*, over 10,000 time steps.

Spiking data was simulated according to a marked point process model with two peaks, each corresponding to a place cell. Both the spike time and mark are generated as a function of the process *x_t_* and the mark can be thought of as a waveform amplitude. The two peaks are centered at 2 and −2 in position and 11 and 12 in mark space. These peaks are each modeled as Gaussian functions with peak values of 0.15 spikes per time step and covariance matrix [0.5, 0; 0, 0.09]. This leads to moderate overlap between the peaks in the mark space (making perfect spike sorting impossible) but minimal overlap in the place coding. Finally, each neuron has a refractory period defined by the negative of a Gaussian function, centered at zero lag after a spike and with a standard deviation of 14 time steps, and neuron 2 has an excitatory influence on neuron 1 defined by a positive Gaussian function, centered at lag 10 time steps after a spike and with a standard deviation of 2 time steps.

Mathematically, the population spiking model is given by the joint mark intensity function

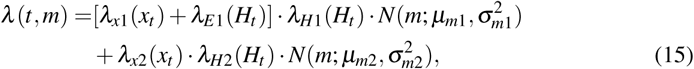

where

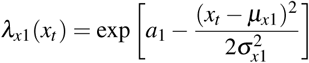

and

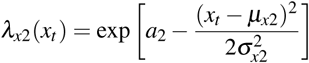

represent the place fields for neurons 1 and 2 respectively,

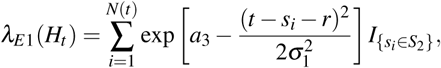

represents the excitatory influence of neuron 2 on neuron 1,

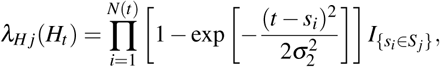

**Table 1.** Simulation study model parameters

| $a_1$ | $a_2$ | $a_3$ | $\mu_{x1}$ | $\mu_{x2}$ | $\sigma_{x1}$ | $\sigma_{x2}$ | $\mu_{m1}$ | $\mu_{m2}$ | $\sigma_{m1}$ | $\sigma_{m2}$ | $\sigma_1$ | $\sigma_2$ | r |
| --- | --- | --- | --- | --- | --- | --- | --- | --- | --- | --- | --- | --- | --- |
| $\log(0.15)$ | $\log(0.15)$ | $\log(0.3)$ | -2 | 2 | $\sqrt{.5}$ | $\sqrt{.5}$ | 11 | 12 | 0.3 | 0.3 | 2 | 14 | 10 |

for *j* = 1, 2 represents the refractoriness of neuron *j*, and 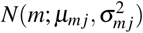 expresses the normal distribution of marks for neuron *j*.

Here, *S*_1_ and *S*_2_ are the sets of spike times from neuron 1 and 2, *a*_1_ and *a*_2_, *μ_x1_* and *μ_x2_*, *μ_m1_* and *μ_m2_*, 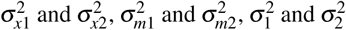 are the numeric values for the peak firing rates, centers in location space and mark space, variance in location space and mark space of these two place cells, variance of excitatory influence and refractoriness respectively, and *N*(*t*) is the total number of spikes up to time *t*. The numeric values for these constants used in the simulation can be found in table 1.

Fig. 1 shows the simulated spiking from this population as a function of the simulated *x_t_* trajectory. In the top panel, spikes are shown as a function of time and position as red and blue dots. The red and blue coloration indicate whether a spike comes from neuron 1 or neuron 2, respectively. We can see a set of red spikes that tend to occur whenever *x_t_* is near −2, and a set of both blue and red spikes that occur whenever *x_t_* is near 2. This is due to the place field of neuron 2 and its excitatory influence on neuron 1. Note that the purpose of this simulation is not to mimic actual place field populations accurately and find the best model to fit, but to generate data that will provide intuition and highlight the ability of the general time-rescaling theorem to assess the goodness-of-fit in data with different types of dependence structures.

Using the simulated data, we performed goodness-of-fit analysis using the time-rescaling theorem we developed above on three possible spiking models. The first uses the true model that generated the data from Eq. 15, including the correct structure for the place fields and the mark distribution, and the full history dependence capturing the refractoriness of each neuron and the excitatory influence of neuron 2 on neuron 1.

The second model uses the correct place and mark structure of the spiking, but omits the history dependent structure completely. Mathematically, this is given by the joint mark intensity function

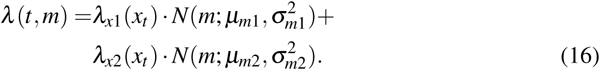

The third model uses a crude spike sorting procedure based on whether each mark value is above or below 11.5, to fit individual intensity models for each of the two sorted neurons. Each neuron has the correct place field structure and history dependent structure, but some spikes are mis-sorted due to the overlap in the mark distribution. Mathematically, the pair of the intensity models for these neurons are given by the following equations:

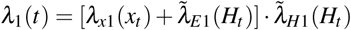

and

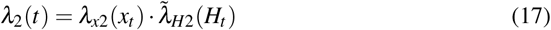

**Fig. 1.**
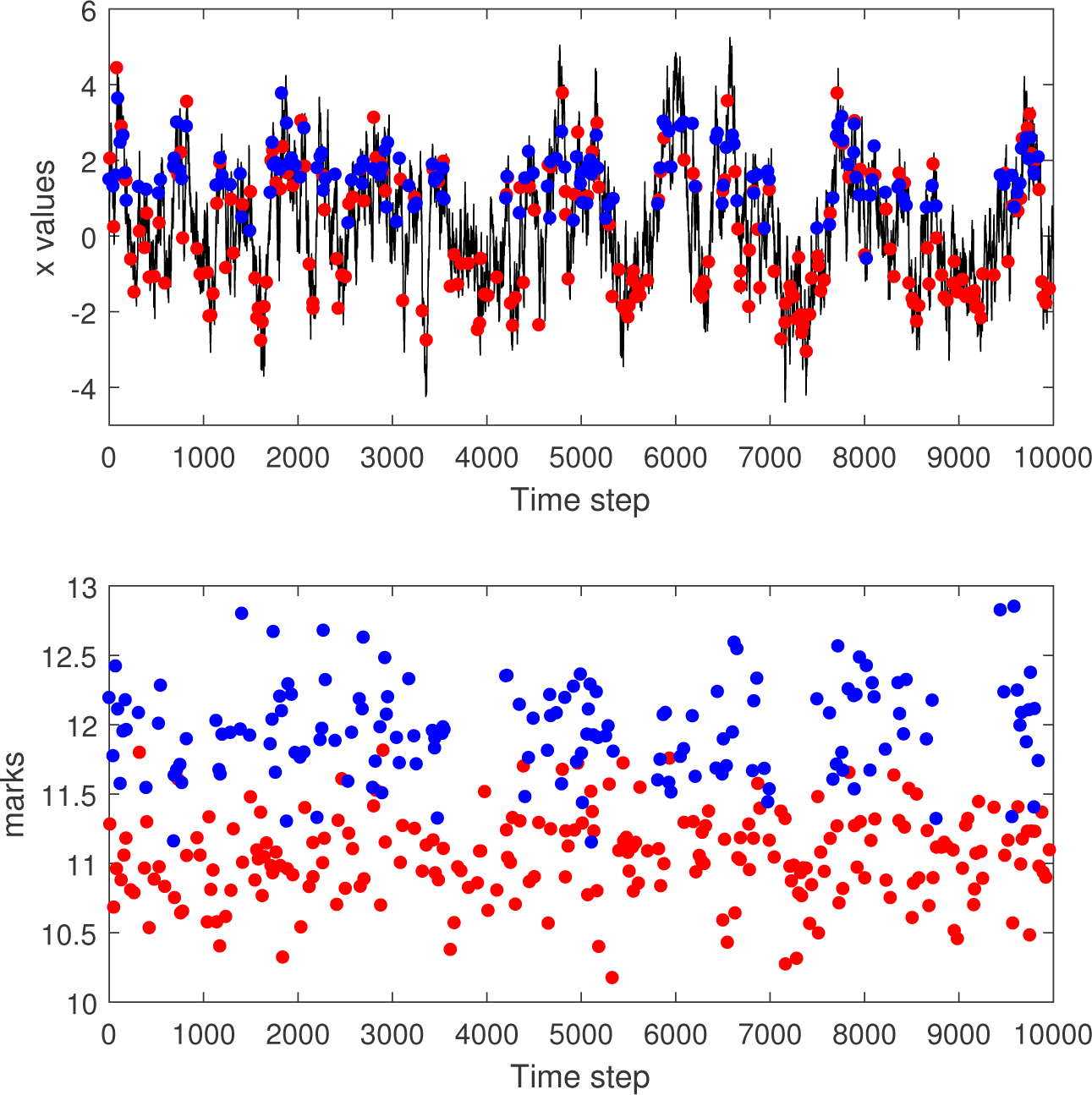
Simulated spiking from a marked point process model with joint mark intensity that depends on a state variable xt defined as an AR(1) process, as defined in Eq. 15. **Top panel:** simulated x-values and spike locations in time. **Bottom panel:** mark values of each spike. Red and blue spike colors indicate whether a spike comes from neuron 1 or neuron 2.

where the excitatory and refractory history dependent component now use the sorted spike identities:

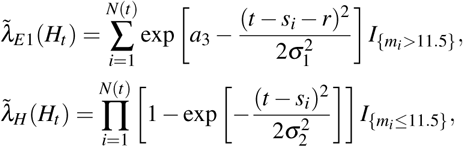

and

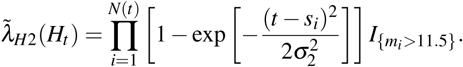

**Fig. 2.**
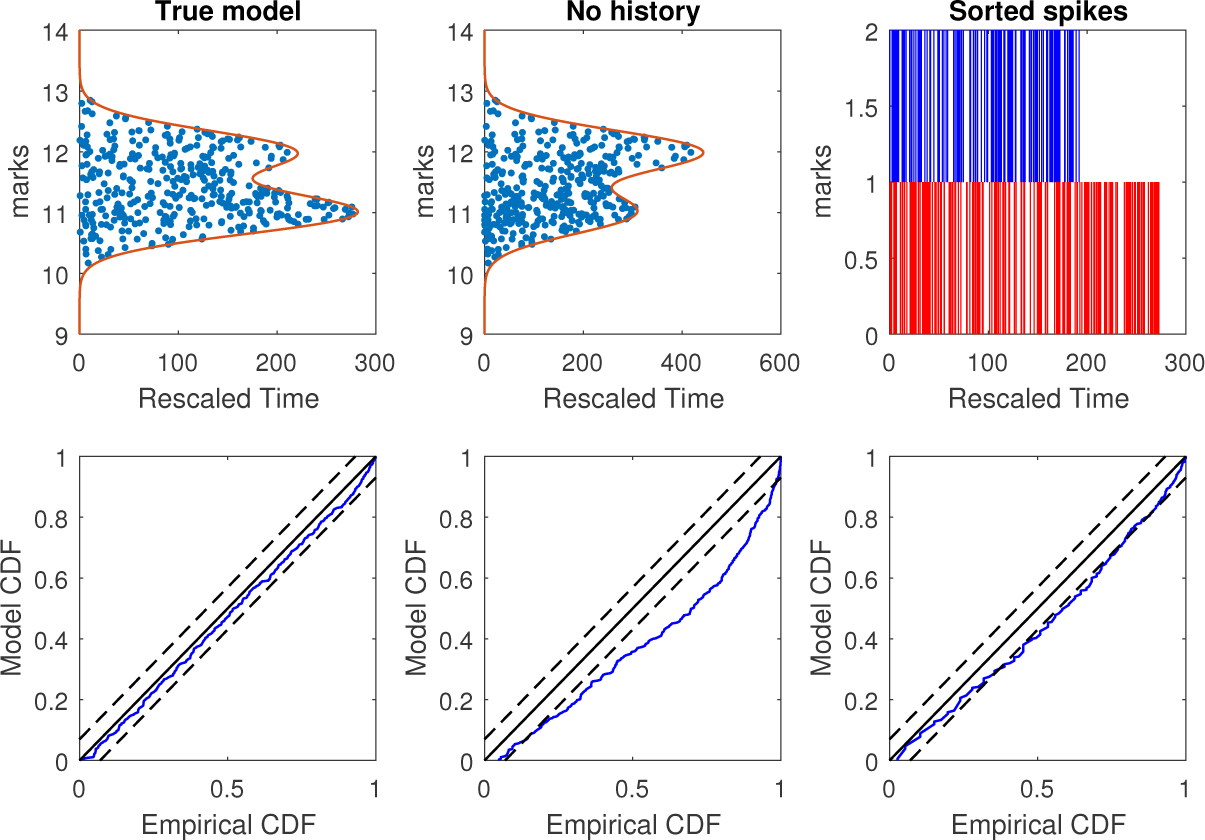
Goodness-of-fit analysis for simulated data based on three candidate models: Left panels use the true model that generated the data, including correct structure for place fields, marks, and history dependence; Middle panels use a model that includes the correct structure for place fields and marks, but omits the history dependence; Right panels use a model that includes correct structure for place fields and history dependence, but uses crude spike sorting rather than true mark structure. Top panels show rescaled spike times (blue dots) and observation intervals (red line) across all mark values. For spike sorted model, rescaled times for each cluster are shown. Bottom panels show KS plots based on all rescaled spike times. The true model produces rescaled spikes that are uniformly distributed in time-mark space with p-value= 0.85 for the Pearson chi-square test and a KS plot that stays within 95% confidence bands. The Pearson chi-square test for the history dependence missing model has p-value=2 *×* 10^−6^, indicating non-uniformity of rescaled spikes. The two intensity models for the sorted spikes demonstrate lack of fit in the KS plots.

Fig. 2 shows the results of the time-rescaling analysis applied to each of the proposed models, using the same simulated spike data. The left panels show the goodness-of-fit assessment based on the true model used to generate the data given by Eq. 15. The middle panels show the goodness-of-fit for the marked point process model in Eq. 16 with correct mark and state dependence, but missing the history dependent component. The right panels show the goodness-of-fit based on crudely sorted spikes given by the models in Eq. 17 with the correct state and history dependence structure. The top panels show the rescaled spike times for each model. For the top-right panel, this is just the rescaled spikes for the two sorted neurons. For the left and middle panels, the rescaled spike times are given by blue dots, and the rescaled values of the end of the observation interval, *τ* (*m,T*), are shown as a function of *m* as a solid red line. The bottom panels show KS plots for all of the rescaled spike times under each of these models.

For the true model, the value of *τ*(*m,T*) has local peaks around mark values of 11 and 12, corresponding to the two peaks in the joint mark-intensity function as these values. The peak around *m* = 11 is larger because of the excitatory influence in the history dependence from neuron 2 to neuron 1. Visually, the rescaled spike times appear to fill out this rescaled time-mark subspace uniformly. A Pearson chi-square test for homogeneity of the rescaled times in this interval yields a p-value of 0.85, suggesting no clear evidence of inhomogeneity. The KS plot everywhere stays within its 95% significance bounds, suggesting no clear lack of fit among the full set of rescaled spike times.

**Fig. 3.**
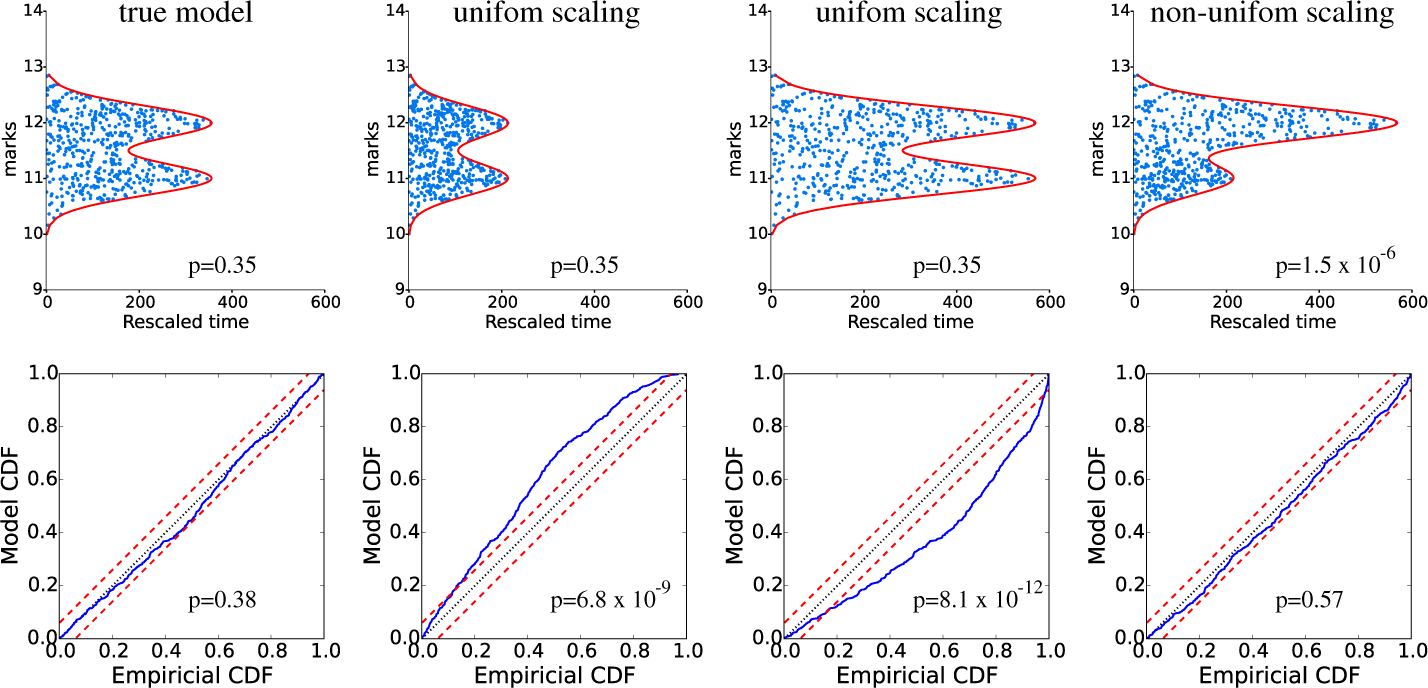
Goodness-of-fit analysis for simulated data based on four candidate models: The top panels show the rescaled spikes (blue) and region, R (red), and the p-values for the chi-square test; the lower panels show KS plots and corresponding p-values. The models are (left panel) true model, (next two panels) true model *λ*(*x_t_*, *m*) scaled uniformly by 0.56 and 1.6, and (right panel) true model whose components *a_c_*, *c* = 1, 2 scaled separately by 0.56 and 1.6, respectively. The last model roughly preserves overall firing rate. KS plots detect the correctness of the firing rate irrespective of mark value, while the Pearson chi-square test characterizes how well the model captures mark structure of the joint mark intensity, which explains why the middle two wrong models still pass the Pearson chi-square test, while the last model passes the KS test.

For the marked point process model missing the history dependent structure (middle panel), the peak around *m* = 12 is larger because *x_t_* stays near the place field of neuron 2 more often than neuron 1, while the missing history dependence does not affect the intensity. By eye, it seems that the rescaled times for mark values below 11.5 occur more densely than those for mark values above 11.5. This is borne out by the Pearson chi-square test (*p* = 2 *×* 10^−6^), which suggests inhomogeneity on the rescaled times, and therefore lack of fit between the model and the original spike data. The lack of fit is also visible in the KS plot, where the observed rescaled interspike intervals are consistently significantly larger than the model estimates.

The panel on the top right shows the rescaled times based on two sorted clusters. As a population model, this could be considered as a marked point process where the marks represent the cluster assignment. In that case, rescaling each spike according to its mark is equivalent to rescaling based on the intensity for whichever neuron the spike is clustered into. Missorted spikes therefore tend to be incorrectly scaled, leading to lack of fit, as observed through the KS plot.

### 3.2 Simulation study 2

We performed a second simulation to illustrate how the KS plot and chi-square test highlight different aspects of the goodness-of-fit. We consider again the same two neurons tuned to a single covariate *x_t_*, and remove the history dependence of spiking, so that two neurons are simply inhomogeneous Poisson spiking units. In this case, the true joint mark intensity model, from which we generate the data, has the same form as Eq. 16, with parameters given in Table 1.

Fig. 3 shows a goodness-of-fit analysis on the resulting data for four different candidate models we propose, (left panel) the true model, (next two panels) two models whose *λ*(*t,m*) are uniformly scaled by 0.56 and 1.6, respectively, and (right panel) a non-uniformly scaled model, with *a_c_* (from Table 1) scaled separately by 0.56 and 1.6, for *c* = 1, 2. Rescaling of the spikes according to the true model *λ*(*t,m*) produces good fits according to both tests, while the uniformly scaled candidate models pass the Pearson chi-square test with p-value 0.35, but the KS plot are far from being in the 95% confidence bounds. The deviation direction from the 45 degree line can be used to determine that the misspecified models underestimate and overestimate the intensity, respectively. The non-uniformly scaled candidate model, where each neuron has been scaled separately while keeping the overall firing rate about the same close to that generated by the true model, passes the KS test, but the chi-square p-value is very small at 1.5 *×* 10^−6^. While the overall pattern of rescaled interspike intervals doesn’t show lack of fit, the rescaled spikes are more concentrated at low mark values and less concentrated at high mark values. This example illustrates the importance of having multiple goodness-of-fit approaches to characterize different features of the data that may be captured or misspecified by a model. In this example both the KS analysis and the Pearson chi-square test are enabled by time-rescaling of the marked point process.

## 4 Data analysis

We analyzed recordings from tetrodes placed in the CA3 region of hippocampus of a rat traversing a W-shaped environment, performing a continuous alternation task. Spikes were detected offline by choosing events whose peak-to-peak amplitudes were above a 40 *µV* threshold in at least one of the channels. For each spike the peak amplitudes across each electrode channel were used as a 4-dimensional mark. Some spikes with lower amplitude peaks may include events whose origin may not be from well-isolated neurons sought in traditional spike-sorting, and may well simply be electrical noise. These spikes are referred to as “hash spikes”, and exist on a continuum extending below the single channel threshold often used for spike detection. In our clusterless population model, we include these hash spikes. We model the joint mark intensity using a mixture of Gaussians

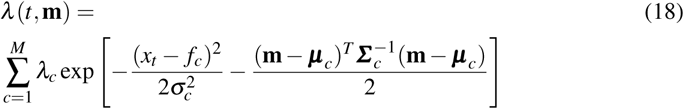

where *M* is the number of Gaussian components, *x_t_* is the position of the animal, m is the four-dimensional mark vector, and *f_c_*, 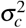 and *μ_c_*, *Σ_c_* are the means and covariances in position and mark spaces, respectively. The parameters were all estimated using a Gibbs sampling procedure (Geman and Geman, 1984; Gelfand and Smith, 1990).

Fig. 4 shows the mark data from a single example tetrode. Fig. 4B shows the time and position of occurrence of unsorted spikes, tracing out the path a rat traveled in the maze. Fig. 4C, right, shows the mark value of each spike, the spike peak amplitude on each channel, as a function of time. Spikes are seen to occur preferentially at certain times, indicating place specific firing from one or more neurons. Fig. 4C, left, shows the 5-dimensional joint mark intensity function, estimated based on Eq. 18, displayed two dimensions at a time. The rightmost column shows the place-specific firing structure, while the rest of the rows show various projections of the spike waveform features familiar to practitioners of manual spike sorting.

Fig. 5 shows the results of a time-rescaling analysis of the joint mark intensity function shown in Fig. 4C. The scatter plot in Fig. 5A shows the corresponding rescaled times and marks for each channel. Here, we do not expect the time-rescaled spikes to appear uniform for each 2D projection, since the other mark dimensions on which the rescaling depends have been collapsed, leading to high density at shorter times. The KS plot in Fig. 5B shows that the rescaled spikes stay in the 95% confidence bounds, suggesting that the model captures the structure of the overall firing rate well. We also performed the Pearson chi-square test by counting the number of spikes occurring within non-overlapping subsets of the 5-dimensional bounded region of the mark space. Dividing the bounded region into subsets where the expected number of spikes is at least 10, we obtained 93 regions for the entire bounded region, giving a p-value of 1.1 *×* 10^−12^, suggesting poor fit of the mark-dependent features. A possible culprit of this is a poor fit is the hash spikes, which require the choice of an arbitrary threshold for the peak-to-peak height, often resulting in a visible cutoff in their distribution. Comparing the spikes to the fitted joint mark intensity function for channel 1 vs. 4 in Fig. 4C, for example, we see where the fitted function clearly does not coincide in the lower amplitudes with the spikes, a consequence of using the mixture of Gaussian intensity model. To investigate this possibility, we performed another Pearson chi-square test, this time only including regions that contained no hash spiking. The resulting p-value of 0.015 still suggests model lack of fit, but a substantial improvement over the full model. In this example, the goodness-of-fit analysis based on time-rescaling has allowed us to characterize features of the data that are explained well by the models, features that are not captured, and how the model might be refined to improve goodness-of-fit.

## 5 Discussion

In this paper, we developed a general toolbox for assessing statistical models of neural populations based on a generalization of the time-rescaling theorem. Given technological advances in neural data acquisition, experiments involving multiple electrodes have now become standard in the practice of neuroscience, making these neural population models of great interest. Understanding these network structures sheds light on how groups of neurons interact with, react and respond to one another and help define possible functions of regions of the brain (Chen et al, 2011; Macke et al, 2011). In addition, the prevalence of multiunit data has brought into question the necessity of spike sorting in every neural population analysis. While many population analyses begin with a spike sorting step and a characterization of the receptive field properties of each sorted neuron, multiple recent experiments have explored the power of clusterless population models (Kloosterman et al, 2014; Deng et al, 2015). Therefore it is valuable to have goodness-of-fit tools that can apply equivalently to both sorted and clusterless population models.

A fundamental challenge in assessing the goodness-of-fit of models of spiking systems is that the timing of each spike has its own distribution, based on many factors that can include coding of dynamic biological and behavioral variables, past spiking history, network effects, and adaptation. The time-rescaling theorem allows us to take any candidate model, and all the dependence structures it describes, and rescale the spikes in such a way that, if the model is correct, they should become samples from a simple uniform distribution. We can then use well-established methods for assessing uniformity to assess the quality of the original model used for rescaling. Furthermore, by taking only the rescaled spike times and disregarding the marks, we can generate a new univariate spike train and use the many existing goodness-of-fit tools for individual spike trains to assess the quality of the joint mark intensity model.

**Fig. 4.**
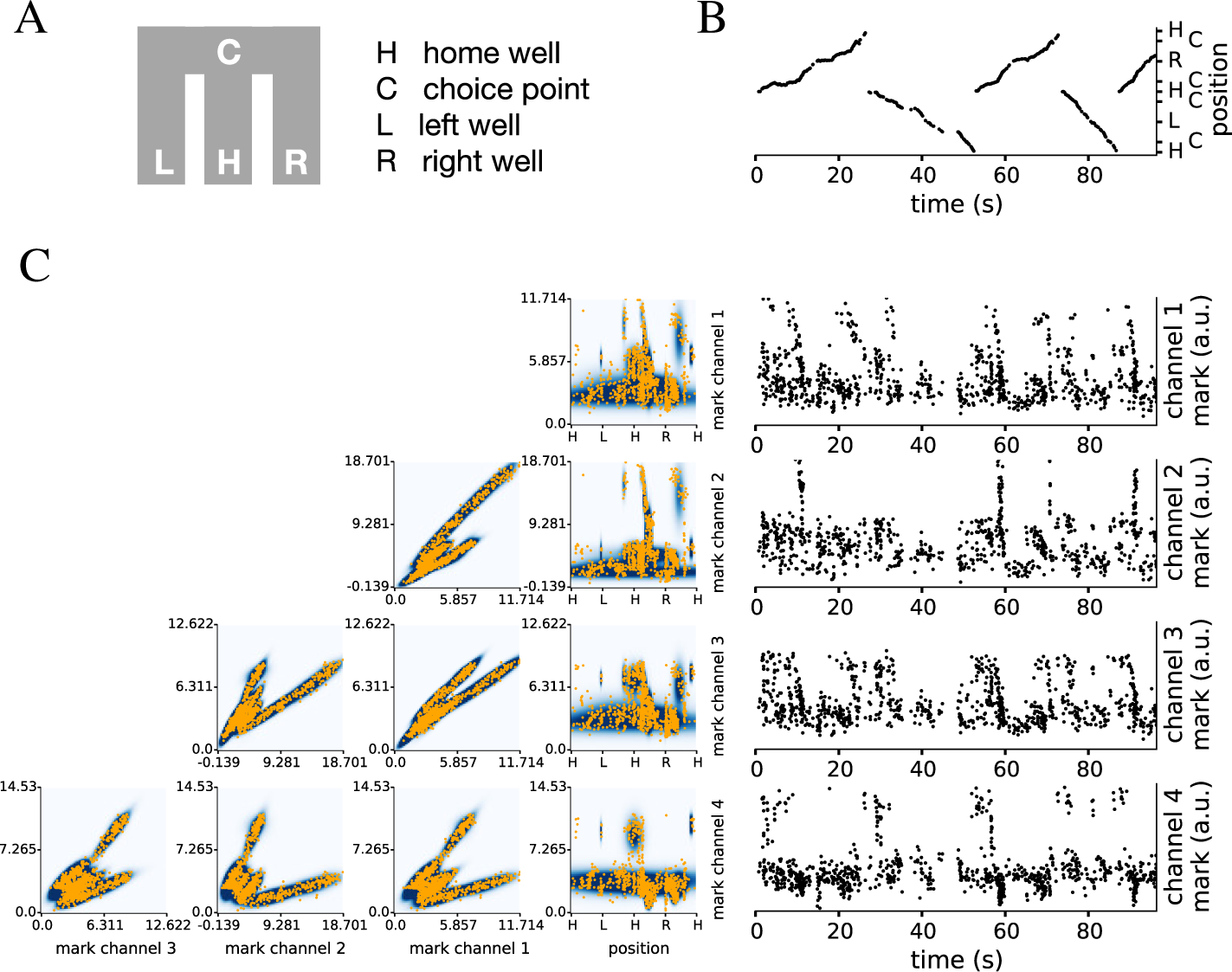
Unsorted spikes and their marks from rat CA3 as it traverses a W-shaped maze, and fitted mark intensity function: A. Schematic of the maze. There are 4 landmarks, the home well H, the choice point C and the left L and right R reward wells. B. Timing and location of all observed spikes in the 1-dimensional position representation. C. Left, the fitted joint mark intensity function and spikes, shown in all combinations of 2-dimensional projections. Orange dots are spikes, and the darker color shows higher intensity. Right, timing and mark (spike amplitude) in each of the tetrode channels of observed spikes. The clustering of spikes in time is a consequence of place-specific firing.

An important feature of this general time-rescaling theorem is that not only are the spike times rescaled, but the observation interval [0, *T*] is also rescaled for each possible mark value. Since a joint mark intensity model can depend on other stochastic processes (such as its own history or the biological and behavioral variables encoded by the population), the intensity is itself a stochastic process, and therefore the rescaled observation region is also stochastic. Therefore, the assessment of uniformity is based both on the rescaled spike times and the rescaled region.

We illustrated our approach via two simulations as well as an application involving place cell spiking activity from the CA3 region of the hippocampus in a rat performing memory guided navigation task on a W-shaped maze. In the first simulation, we implemented the goodness-of-fit tests in three different model fits: the true model, a model intentionally missing history-dependence, and a model for which the mark corresponds to one of two labels given by spike sorting. As expected, the results indicated proper fit with the true model, and a lack-of-fit in both the model missing history dependence as well as the sorted model. This demonstrated the ability of the approach to discern different reasons for lack of fit. Importantly, we could assess the quality of fit for both sorted and clusterless spiking models and determine the degree to which sorting affected the model fit. In our second simulation, we demonstrated that distinct goodness-of-fit measures, both based on the same time-rescaling approach, could be used to determine different aspects of the model fit quality. Incorrectly scaling the intensity uniformly over all marks led to lack of fit evidenced by the KS plot but not the assessment of uniformity; differentially scaling subsets of mark values, as might occur with a model that misspecified the receptive fields of particular neurons, led to lack of fit evidenced by lack of uniformity.

**Fig. 5.**
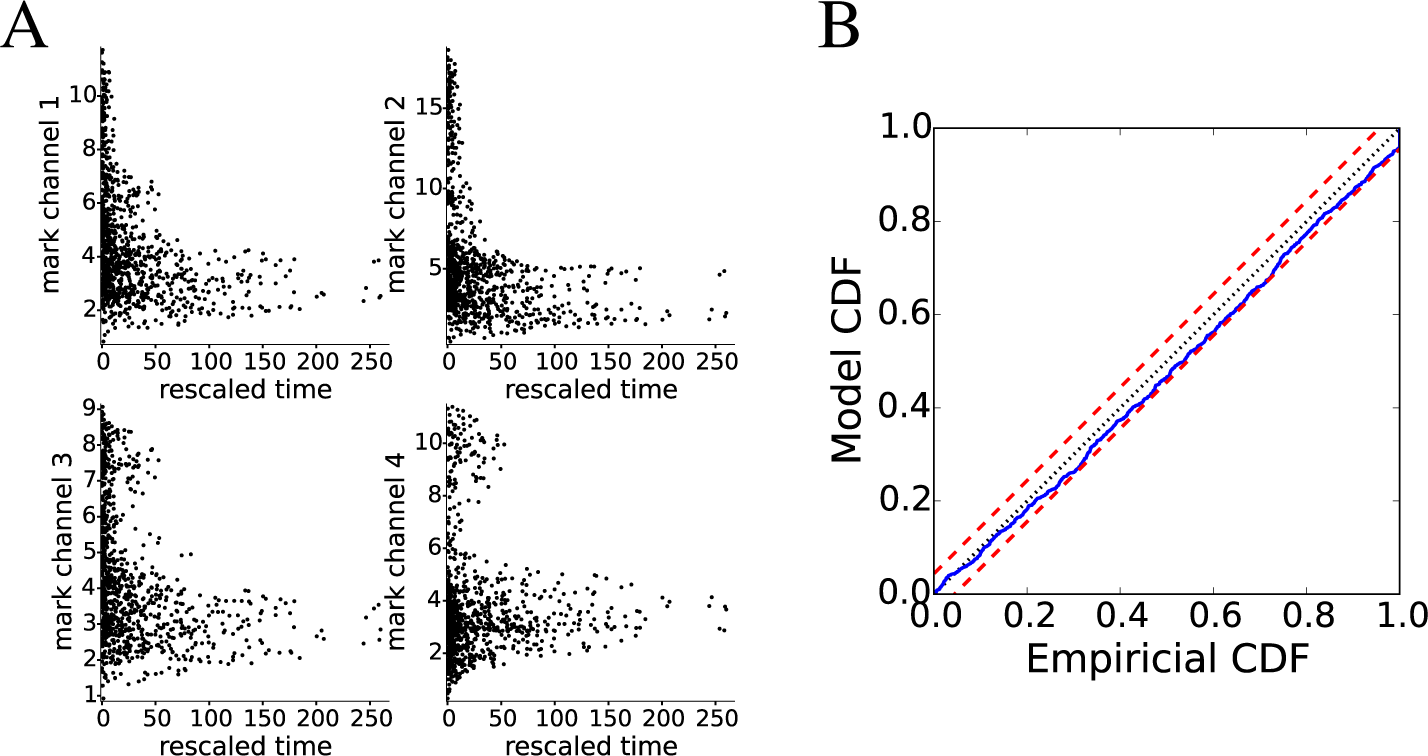
Rescaling analysis for unsorted spikes and their marks from rat CA3: A. Time-rescaled spikes in each of the 4 tetrode channels. The same spikes appear in each of the 4 panels at the same rescaled time, but at different mark values. B The corresponding KS plot.

In our real-data example, we used a 4-dimensional mark representing the waveform peak amplitudes across a tetrode to exhibit the ability to generalize to more complicated mark spaces. The fit of a Gaussian mixture model with no history dependence captured much of the temporal structure, as evidenced by the KS plot, but perhaps fit the spikes in some mark regions better than others, as suggested by the analysis of uniformity. A follow-up analysis, suggested that the model may not be capturing the hash spikes as well as the higher amplitude spikes.

In this paper we focused on two goodness-of fit measures that could be applied to the rescaled spike times and marks, a Pearson chi-square test comparing the expected and observed number of spikes in subsets of the rescaled observation region, and a KS plot analysis based on rescaling the rescaled spike times again based on the expected rescaled spiking rate. However, there are a variety of other goodness-of-fit tools available after rescaling that could also be used, either instead of, or to compliment these analyses. For example, a well-studied statistical approach for assessing uniformity is based on Ripley’s *K*-function (Ripley, 1977). This function, *K*(*x,r*), is defined as the expected number of points within a ball *b*(*r*) with radius *r* centered at *x*. For uniform rescaled spikes, this function should grow as *r^d^*, where *d* is the dimension of the mark-time space. We can compute the empirical *K* function, 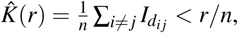, where *d_ij_* is the Euclidean distance between rescaled spikes *i* and *j*, and *I_d_ij__* < *r* is equal to 1 if that distance is less than *r*, and otherwise 0. We can then compare the empirical *K* function to the theoretical one under a uniform model to assess the quality of our original model. We can also construct confidence intervals for the estimated function and compute a corresponding p-value via Monte Carlo simulations (Baddeley et al, 2005). A variety of other well-documented and tested methods are also available (Petrie and Willemain, 2013) and could be used interchangeably with those we specifically mention in this paper. A few examples include those that perform a two-sample test on a subsample of points in a high-density region and a subsample in a low-density region (Jain et al, 2002), or those that consider the distribution of distances from points to the boundary of support, both in the case of known support (Berrendero et al, 2006) and unknown support (Berrendero et al, 2012).

There are a number of extensions and avenues for future exploration for this goodness-of-fit framework. In the simulations, we provided examples of how assessments based on time-rescaling could be used to help identify areas of lack of fit, and to suggest refinements to population spiking models. The ways in which different measures might be used for model refinement should be explored in more detail, and specific recommendations could be made about the best measures to use to identify particular features that should be added or altered in a model. Also, in our examples, we limit the standard point process goodness-offit analysis to KS plots but with the appropriate adaptations and generalizations, one could also employ other common techniques such as the QQ plot, autocorrelations of rescaled wait times, or a Fano Factor analysis to assess dispersion.

Another possible extension might focus on mark rescaling rather than time-rescaling. We could retain the observed times of each spike, and modify each spike mark to produce uniform spikes over a stochastic region with a fixed temporal extent, but random mark boundaries. For a one-dimensional mark, this could be achieved by replacing the mark of the *i*th spike, *m_i_*, with the integral 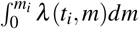 (Merzbach and Nualart, 1986). An advantage of such an approach would be that the spike times would remain the same and be interpretable. For example a cluster of points at a particular time point might suggest model lack of fit specific to that time. However, since the temporal pattern of spikes would be unchanged, it would still retain all the temporal dependence structure in the original data. Additionally, how to best rescale in general mark spaces is still unknown.

Additional research could also be done on improving the computational burden of these methods in high dimensional mark spaces. While the rescaling of times is based only on the number of spikes, not the dimensionality of the marks, the computation of the boundary of the stochastic region will grow in complexity with the mark dimension. There may be multiple ways to deal with this, including methods of efficiently approximating the boundary assuming smoothness of the intensity, or goodness-of-fit measures that are less sensitive or do not require direct knowledge of the full boundary.

With this method, we provide model assessment tools that can be used appropriately for both population models and sorted models and and help collect more detailed information on their respective fits. In this way, researchers can better understand the advantages and disadvantages posed by population and single-unit modeling. Ultimately, this could provide significant insights into the question of when neural network structures can be better understood with spike sorting or direct ensemble modeling. Additionally, as experiments head in more complex directions and datasets become richer, modeling methods will need to improve and develop alongside them. For researchers to maintain confidence in any conclusions drawn from the application of a particular modeling approach, a corresponding goodness-of-fit toolset is essential. Here, we present a general goodness-of-fit approach that can assess and indicate areas of lack-of-fit for a wide variety of population spiking models, enabling researchers to gain more understanding and insight into the increasingly complex data structures being made available in neuroscience.

## Acknowledgements

This work was supported by grants from the NIH (MH105174, NS094288) and the Simons Foundation (542971).

## References

Arai K, Kass RE (2017) Inferring oscillatory modulation in neural spike trains. PLoS computational biology 13(10):e1005,596

Ba D, Temereanca S, Brown EN (2014) Algorithms for the analysis of ensemble neural spiking activity using simultaneous-event multivariate point-process models. Frontiers in computational neuroscience 8

Baddeley A, Turner R, et al (2005) Spatstat: an r package for analyzing spatial point patterns. Journal of statistical software 12(6):1–42

Berrendero JR, Cuevas A, Vjosázquez-grande F (2006) Testing multivariate uniformity: The distance-to-boundary method. Canadian Journal of Statistics 34(4):693–707

Berrendero JR, Cuevas A, Pateiro-López B (2012) A multivariate uniformity test for the case of unknown support. Statistics and Computing 22(1):259–271

Brockwell AE, Rojas AL, Kass R (2004) Recursive bayesian decoding of motor cortical signals by particle filtering. Journal of Neurophysiology 91(4):1899–1907

Brown EN, Frank LM, Tang D, Quirk MC, Wilson MA (1998) A statistical paradigm for neural spike train decoding applied to position prediction from ensemble firing patterns of rat hippocampal place cells. Journal of Neuroscience 18(18):7411–7425

Brown EN, Barbieri R, Ventura V, Kass RE, Frank LM (2002) The time-rescaling theorem and its application to neural spike train data analysis. Neural computation 14(2):325–346

Brown EN, Kass RE, Mitra PP (2004) Multiple neural spike train data analysis: state-of-theart and future challenges. Nature neuroscience 7(5):456–461

Brown TC, Nair MG (1988) A simple proof of the multivariate random time change theorem for point processes. Journal of Applied Probability 25(1):210–214

Chen Z, Putrino DF, Ghosh S, Barbieri R, Brown EN (2011) Statistical inference for assessing functional connectivity of neuronal ensembles with sparse spiking data. IEEE transactions on neural systems and rehabilitation engineering 19(2):121–135

Daley DJ, Vere-Jones D (2003) An introduction to the theory of point processes. Springer, New York

Deng X, Eskandar EN, Eden UT (2013) A point process approach to identifying and tracking transitions in neural spiking dynamics in the subthalamic nucleus of parkinson’s patients. Chaos: An Interdisciplinary Journal of Nonlinear Science 23(4):046,102

Deng X, Liu DF, Kay K, Frank LM, Eden UT (2015) Clusterless decoding of position from multiunit activity using a marked point process filter. Neural computation

Eden UT, Frank LM, Barbieri R, Solo V, Brown EN (2004) Dynamic analysis of neural encoding by point process adaptive filtering. Neural computation 16(5):971–998

Eden UT, Frank LM, Long T (2018) Characterizing Complex, Multi-Scale Neural Phenomena Using State-Space Models. In: Dyn. Neurosci.

Gelfand AE, Smith AF (1990) Sampling-based approaches to calculating marginal densities. Journal of the American statistical association 85(410):398–409

Geman S, Geman D (1984) Stochastic relaxation, gibbs distributions, and the bayesian restoration of images. IEEE Transactions on pattern analysis and machine intelligence (6):721–741

Georgopoulos AP, Schwartz AB, Kettner RE (1986) Neuronal population coding of movement direction. Science pp 1416–1419

Gerhard F, Haslinger R, Pipa G (2011) Applying the multivariate time-rescaling theorem to neural population models. Neural computation 23(6):1452–1483

Huang Y, Brandon MP, Griffin AL, Hasselmo ME, Eden UT (2009) Decoding movement trajectories through a t-maze using point process filters applied to place field data from rat hippocampal region ca1. Neural computation 21(12):3305–3334

Jain AK, Xu X, Ho TK, Xiao F (2002) Uniformity testing using minimal spanning tree. In: Pattern Recognition, 2002. Proceedings. 16th International Conference on, IEEE, vol 4, pp 281–284

Johnson N, Kotz S (1970) Distribution in Statistics-Continuous Univariate Distribution-1 Wiley

Kass RE, Ventura V (2001) A spike-train probability model. Neural computation 13(8):1713–1720

Kass RE, Ventura V, Brown EN (2005) Statistical issues in the analysis of neuronal data. Journal of neurophysiology 94(1):8–25

Kass RE, Eden UT, Brown EN (2014) Analysis of neural data, vol 491. Springer

Kloosterman F, Layton SP, Chen Z, Wilson MA (2014) Bayesian decoding using unsorted spikes in the rat hippocampus. Journal of neurophysiology 111(1):217–227

Macke JH, Buesing L, Cunningham JP, Byron MY, Shenoy KV, Sahani M (2011) Empirical models of spiking in neural populations. In: Advances in neural information processing systems, pp 1350–1358

Merzbach E, Nualart D (1986) A Characterization of the Spatial Poisson Process and Changing Time. Ann Probab 14(4):1380–1390

Meyer PA (1971) Démonstration simplifiée d’un théorème de knight

Paninski L, Pillow J, Lewi J (2007) Statistical models for neural encoding, decoding, and optimal stimulus design. Progress in brain research 165:493–507

Paninski L, Brown EN, Iyengar S, Kass RE (2009) Statistical models of spike trains. Stochastic methods in neuroscience pp 278–303

Papangelou F (1972) Integrability of expected increments of point processes and a related random change of scale. Transactions of the American Mathematical Society 165:483–506

Petrie A, Willemain TR (2013) An empirical study of tests for uniformity in multidimensional data. Computational Statistics & Data Analysis 64:253–268

Pillow JW, Shlens J, Paninski L, Sher A, Litke AM, Chichilnisky E, Simoncelli EP (2008) Spatio-temporal correlations and visual signalling in a complete neuronal population. Nature 454(7207):995–999

Port SC (1994) Theoretical probability for applications, vol 206. Wiley-Interscience

Ripley BD (1977) Modelling spatial patterns. Journal of the Royal Statistical Society Series B (Methodological) pp 172–212

Ross SM (1996) Stochastic processes. 1996. Wiley, New York

Shanechi MM, Hu RC, Powers M, Wornell GW, Brown EN, Williams ZM (2012) Neural population partitioning and a concurrent brain-machine interface for sequential motor function. Nature neuroscience 15(12):1715–1722

Smith AC, Brown EN (2003) Estimating a state-space model from point process observations. Neural Computation 15(5):965–991

Sodkomkham D, Ciliberti D, Wilson MA, Fukui Ki, Moriyama K, Numao M, Kloosterman F (2016) Kernel density compression for real-time bayesian encoding/decoding of unsorted hippocampal spikes. Knowledge-Based Systems 94:1–12

Srinivasan L, Eden UT, Willsky AS, Brown EN (2006) A state-space analysis for reconstruction of goal-directed movements using neural signals. Neural computation 18(10):2465–2494

Truccolo W, Eden UT, Fellows MR, Donoghue JP, Brown EN (2005) A point process framework for relating neural spiking activity to spiking history, neural ensemble, and extrinsic covariate effects. Journal of neurophysiology 93(2):1074–1089

Vere-Jones D, Schoenberg FP (2004) Rescaling marked point processes. Australian & New Zealand Journal of Statistics 46(1):133–143

Wu S, Amari Si, Nakahara H (2002) Population coding and decoding in a neural field: a computational study. Neural Computation 14(5):999–1026

